# EVA: a Foundation Model Advancing Translational Drug Development in Immuno-Inflammation

**DOI:** 10.1101/2025.05.02.651839

**Authors:** Aziz Fouché, Apolline Bruley, Matthew Corney, Pierre Marschall, Vincent Bouget, Julien Duquesne

## Abstract

Drug development is a lengthy and high-risk process, with most investigational drug candidates failing in phase II randomized clinical trials (RCT) due to insufficient efficacy. It makes early prediction of trial outcomes crucial for reducing attrition and guiding strategic decisions, especially in immunology and inflammation (I&I) diseases. Herein, we present EVA, the first pre-trained foundation model in complex inflammatory diseases tailored to support drug development. EVA learns generalizable patterns from large-scale data of cell biology and immunology, enabling superior predictive performance and generalization compared to traditional approaches. EVA is pre-trained on tens of millions of single-cell RNA-seq samples and tens of thousands of bulk RNA-seq samples from I&I diseases patients, enabling it to learn disease-relevant transcriptomic patterns in this therapeutic area. By fine-tuning EVA in few-shot settings on both preclinical (mouse) and clinical (human) data and harnessing its wide pre-training knowledge, EVA predicts drug responses in I&I with high precision at both cohort and patient levels, as illustrated by accurate forecasting of anti-TNF therapeutic activity in ulcerative colitis. By deciphering its decision process, we further highlight that EVA’s ability to stratify patients based on predicted drug response can also be leveraged to discover drug response biomarkers as early as preclinical stages. EVA’s applications in precision immunology encompass therapeutic target validation prior to clinical entry, identification of patient subpopulations most likely to benefit from treatment, and comparative efficacy analysis against competitor compounds. EVA’s versatility makes it an invaluable tool for strategic decision-making throughout the drug development pipeline: by leveraging it to prioritize the most promising drug candidates and optimize RCT designs, it can contribute to reduce late-stage failures and accelerate the delivery of effective therapies. Overall, this work represents a significant advancement in utilizing a pre-trained foundation model for precision drug development in complex inflammatory diseases.

**Graphical abstract:** EVA is a pre-trained foundation model specific to immune-mediated inflammatory diseases. It enables the prediction of therapeutic efficacy in patients leveraging data from preclinical disease models.

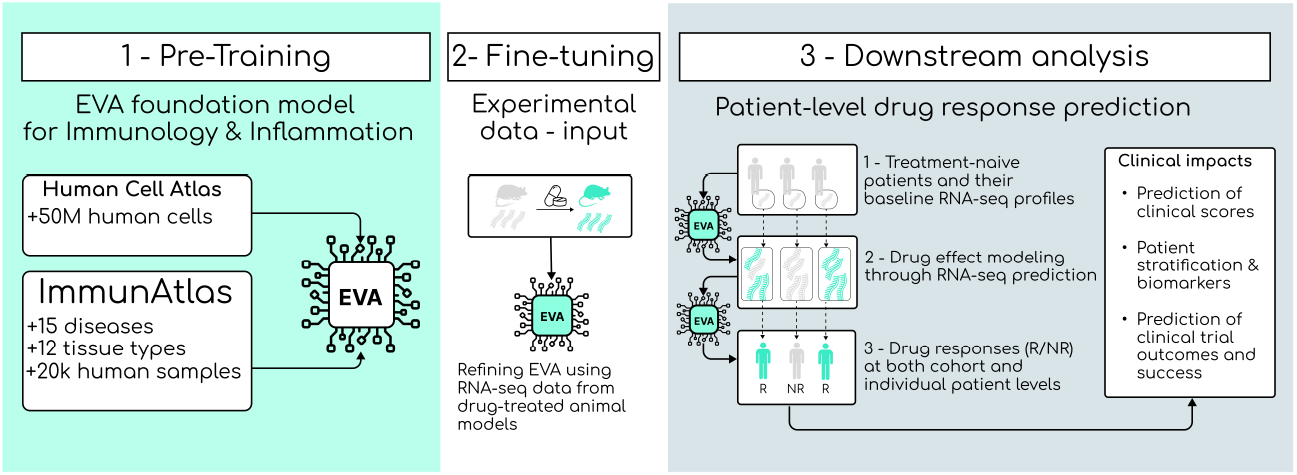

## Introduction

Immunology and inflammation (I&I) diseases constitute a diverse group of chronic conditions characterized by a dysregulated immune system, leading to sustained inflammation and tissue damage in target organs. I&I diseases affect approximately 5–7% of the population in industrialized countries and show a rising prevalence in South America and Asia^1^,^2^. These disorders encompass rheumatological, dermatological, respiratory, and gastrointestinal affections, such as inflammatory bowel diseases (IBDs), including Crohn’s disease and ulcerative colitis (UC). The societal impact of these chronic conditions is immense, with substantial economic consequences through direct healthcare costs exceeding billions annually, reduced workforce productivity, significantly diminished quality of life for patients who often face lifelong disability and psychological distress, and increased rates of malignancy due to chronic inflammation and immunosuppressive treatments. Despite the high burden of these diseases, the success rate in developing new drugs for immunology and inflammation conditions remains low, with a failure rate of up to 95%^3^.

The broad range of manifestations observed in I&I diseases is increasingly understood as reflecting a shared disease spectrum rather than isolated, organ-specific conditions. This shifting paradigm is bolstered by convergent evidence from various domains: common underlying molecular and immune mechanisms, frequent clinical co-occurrence across diseases, overlapping symptomatology, and familial clustering. These clinical patterns are underpinned by shared genetic susceptibility loci and immunopathogenic pathways— particularly those involving dysregulation of specific cytokines including TNF-α, IL-17, and IL-23^4-6^. Nevertheless, despite these shared features, a subset of patients—whose size varies depending on the disease—experiences suboptimal treatment outcomes or relapses after an initial period of improvement^7,8^. This reflects an incomplete understanding of the underlying pathophysiological mechanisms in I&I diseases, which limits the development of optimal patient stratification strategies to maximize therapeutic response and hinders clinical trial success.

Furthermore, the clinical heterogeneity of I&I diseases, often presenting with overlapping symptoms and organ involvement, necessitates the use of composite disease activity scores and multifaceted endpoints in clinical trials^9^. This complexity introduces hurdles that make study design more arduous and further diminishes the probability of positive outcomes. These obstacles likely contribute to the high attrition rate observed in phase II clinical trials—often referred to as the “valley of death”—where promising drug candidates frequently fail to demonstrate sufficient efficacy in the clinic^10^.

Translational research, which bridges the gap between basic research and clinical trials, is a critical stage in drug discovery & development. Pivotal decisions made at this stage— typically based on limited preclinical and early clinical data—can determine a drug candidate’s future success. These include the design of the clinical development plan, identification of companion biomarkers, and strategic positioning within the competitive landscape. Because these decisions are only validated through costly Phase II and III trials, thoughtful and rigorous trial design is essential.

Preclinical models such as cell lines, animal disease models, and organoids are indispensable for generating early-stage data informing these choices. Yet these models have significant limitations, as they rarely replicate human physiology or capture the complexity of systemic diseases like I&I diseases. While human clinical trials are expensive and time-intensive, preclinical data is more readily available and less costly to generate. Coupled with recent breakthroughs in foundation models, this offers a promising avenue to mitigate the economic burden of drug development by enabling earlier and more accurate *in silico* identification of high-potential candidates.

Modelling the effect of a drug is an important challenge in computational pharmacology, opening opportunities to predict the outcomes of clinical trials, prioritize or repurpose existing compounds, and discover new therapeutic targets. Various modeling approaches have been elaborated over the years, typically based on gene regulatory networks (GRN) and quantitative systems pharmacology (QSP)^11,12^. Unfortunately, developing these computational models requires extensive literature review, trial-and-error, and expert fine-tuning, all of which must be repeated for each new drug or disease. This limited scalability prevents their generalized adoption in industrial settings, where hundreds of potential drug-disease models would need to be developed. Furthermore, these models oversimply biological systems and are often restricted to literature-reported interactions, limiting their ability to generate new knowledge and introducing strong assumptions whose consequences are difficult to measure. Generalization outside of training data in biology is particularly challenging due to the complex interplay of molecular, cellular, and environmental factors, high dimensionality, noisiness and sparsity of biological data, significant heterogeneity across individuals, and the often-non-linear nature of biological systems. To address this issue, we adopt an innovative approach that leverages a large pre-trained foundation model of I&I to develop a data-driven pipeline that predicts the effects of candidate treatments in I&I diseases.

Foundation models are large-scale pre-trained neural networks that have emerged as a transformative force in canonical machine learning fields, such as natural language processing and computer vision, as well as in computational biology. They already span many biological modalities (DNA, RNA, proteins, histology, clinical data…)^13-19^, and offer unprecedented opportunities in drug discovery and biological research. Ideally, foundation models should be able to leverage the data processed during pre-training to enhance downstream applications, accelerating training, improving generalization even with limited training data, and ultimately delivering more robust and scalable predictions. In practice and especially in computational biology, the current challenges faced by foundation models are (a) the sheer amount of data and computational resources needed during pre-training and (b) their lack of generalization in out-of-domain settings make their application to biological tasks challenging^20,21^, while it is supposed to be the strength of large pre-trained models. Given the abundance of publicly available omics data for pre-training and the shared pathophysiological mechanisms across the spectrum of I&I diseases, we believe there is a strong technical, biological and clinical rationale to develop a foundation model specifically tailored to these diseases.

In this work, we introduce **EVA**, a 50M-parameters foundation model designed for I&I diseases. We demonstrate EVA’s capabilities by predicting the therapeutic effect of anti-TNF for ulcerative colitis patients, using a fine-tuned version of EVA trained with few-shot learning on a small preclinical dataset from another species. Within this integrated framework, we demonstrate EVA’s ability to bridge the gap between preclinical research and clinical impact, offering valuable insights into drug efficacy in humans. Its capacity to forecast the outcomes of a Phase II randomized controlled trial (RCT) using low throughput preclinical data offers a strategic advantage in trial design and resource allocation. By leveraging these predictive capabilities, EVA can significantly enhance the design of clinical trials, thereby improving the odds of success and accelerating the development of effective therapies for I&I diseases.

## Results

### EVA is a foundation model pre-trained on an RNA-seq I&I atlas

Research in the NLP field has underscored the critical role of pre-training and fine-tuning foundation models in enabling their use in practical scenarios^22,23^. While self-supervised pre-training excels at capturing associative knowledge and memorizing patterns that can be retrieved during downstream tasks, raw pre-trained models are often difficult to use effectively without further refinement. Recently, there has been a surge in generalist RNA-seq foundation models for computational biology, but several articles have since highlighted their limitations when applied to real-life use cases^20,21^, as these models still struggle to generalize outside of their training domain. To address this issue, EVA was pre-trained using masked language modeling in a two-step self-supervision process:

∘ We started from a single-cell RNA-seq foundation model based on the transformer architecture that had been pre-trained on a vast corpus of single-cell RNA-seq data, consisting of approximately 50 million human cells of various cell types and tissues gathered from the Human Cell Atlas (HCA)^24^ and the CellXGene portal.
∘ A second pre-training procedure was then conducted using **ImmunAtlas**, a large in-house atlas aggregating public datasets containing in total 21,686 bulk RNA-seq samples of patients suffering from I&I diseases, spanning 15 diseases and more than 10 tissues (Fig. 1a-d). Although ImmunAtlas is significantly smaller than the HCA dataset, it is far more representative of real-world I&I data and contains rich, disease-relevant biologic signals.

**Figure 1:**
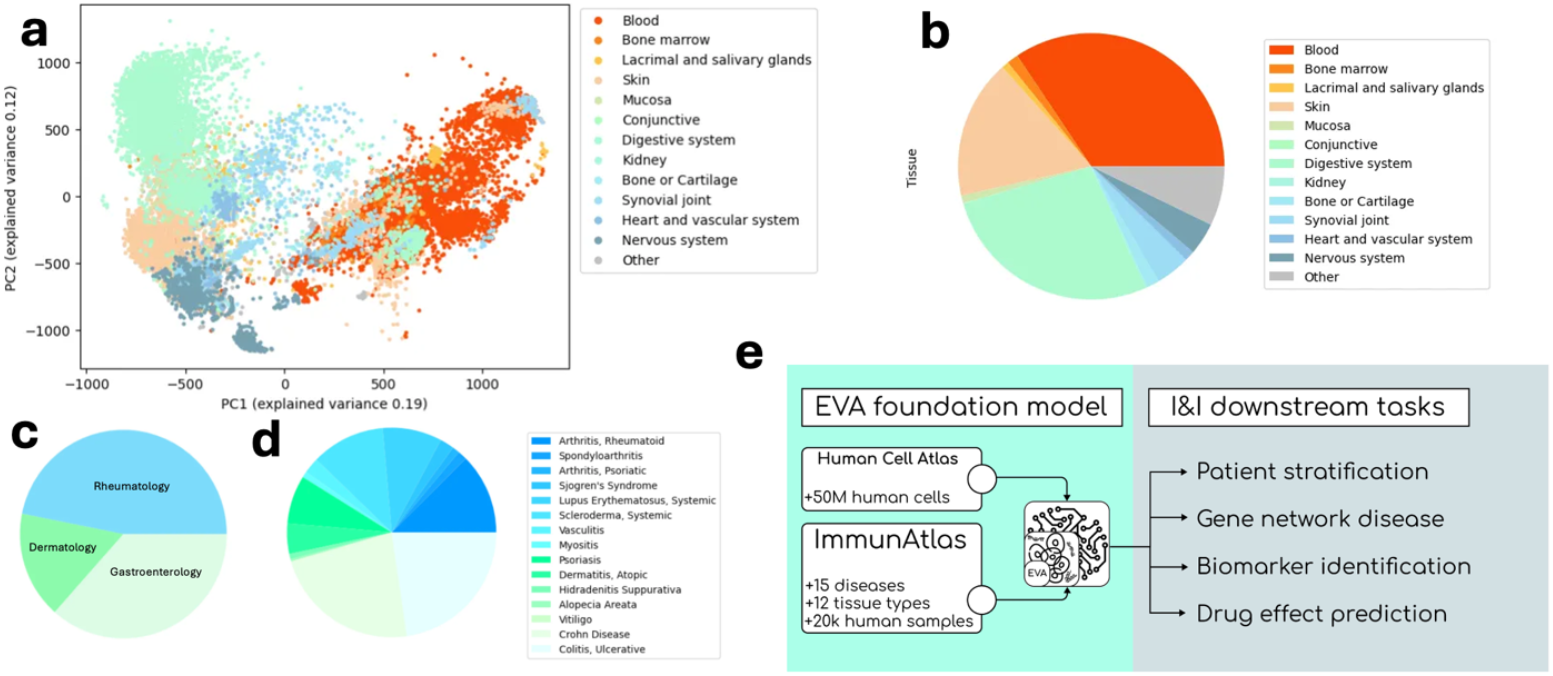
EVA is a foundation model pre-trained on ImmunAtlas, a specific I&I diseases dataset. **(a)** PCA view of the ImmunAtlas dataset, colored by tissue. **(b-d)** ImmunAtlas tissue and I&I disease distribution. **(e)** EVA is an I&I foundation model pre-trained on a large corpus of single-cell and bulk RNA-seq data that can be applied in various I&I downstream tasks.

This training process makes EVA both more efficient when processing bulk RNA-seq data which is still prevalent in the clinic, and more actionable in the context of I&I diseases thanks to its refined representations of immune pathways and gene expression deregulations commonly found in these diseases. If this pre-training step requires a large amount of data and extensive computational power, it must be carried out only once to build the base EVA model. Then, because of its training procedure tailored to I&I and the use of in-house data, EVA can be fine-tuned for various use-cases, requiring much smaller amounts of data and compute which makes it very versatile and scalable. Examples of such downstream tasks that EVA can carry out are disease stratification for biomarker discovery^25^, gene regulatory network inference^26^, or drug response prediction as presented in this paper (Fig 1e). Overall, EVA represents a significant advancement in enhancing the utility and precision of foundation models for biomedical research in I&I.

### EVA learns in few-shot the effect of TNFR2 KO from mouse preclinical data

A significant milestone in drug effect modeling is the ability to accurately predicting transcriptome-wide gene expression changes in response to perturbations, such as gene knock-out or drug administration. We hypothesize that modeling the transcriptomic shift in response to a drug in the primary organ affected by a disease can be used to forecast the compound’s therapeutic impact at the phenotypic level. In recent years, several approaches have explored the in silico prediction of drug-induced transcriptomic changes^13,27^; however, these models often struggle to outperform basic baselines, such as naive predictors that simply estimate the mean expression change across samples^20^. In preclinical settings in particular, the complexity lies in (a) the scarcity of preclinical data, typically limited to 5-10 individuals from a model organism (such as mice or cell lines) per drug candidate, requiring models that can generalize from very limited training data; and (b) the difficulty in translating findings from model organisms to humans, causing potential out-of-distribution issues. Thanks to its pre-training knowledge of I&I coming from ImmunAtlas, EVA is uniquely positioned to address these two challenges. It can leverage its learned representations of gene associations, biological pathways, and human biology of the diseases of interest to better project a compound’s impact on human subjects, even with minimal available data.

To demonstrate its potential, we evaluated EVA on the following perturbation task: inferring transcriptome-wide changes resulting from TNF receptor 2 (TNFR2) depletion in mouse, using a small case-control dataset from an IBD mouse models. The perturbation dataset consisted of six mice with IBD, including three control individuals and three which have got their TNFR2 gene knocked out (TNFR2-KO)^28^, mimicking the effect of an anti-TNF treatment. Because mice were from an inbred line, each TNFR2-KO mouse could be paired with each TNFR2-sufficient one, thus generating 9 matched case-control pairs (95k gene expression differences for training, 27k left out for validation). We employed a stochastic encoder-decoder architecture to learn the TNFR2-KO-perturbation for each input gene as the parameters of a multivariate Gaussian (MVG) probability distribution with diagonal covariance, comparing EVA to several state-of-the-art RNA-seq foundation model for the encoder part. We also established a basic MVG with diagonal covariance, fitted to the training data, as a baseline for comparison. We fine-tuned all foundation models on the same training dataset and evaluated their performance on the validation set over 10 random initializations, using the average Spearman’s correlation between expected and predicted perturbations as the metric (Fig. 2).

**Figure 2.**
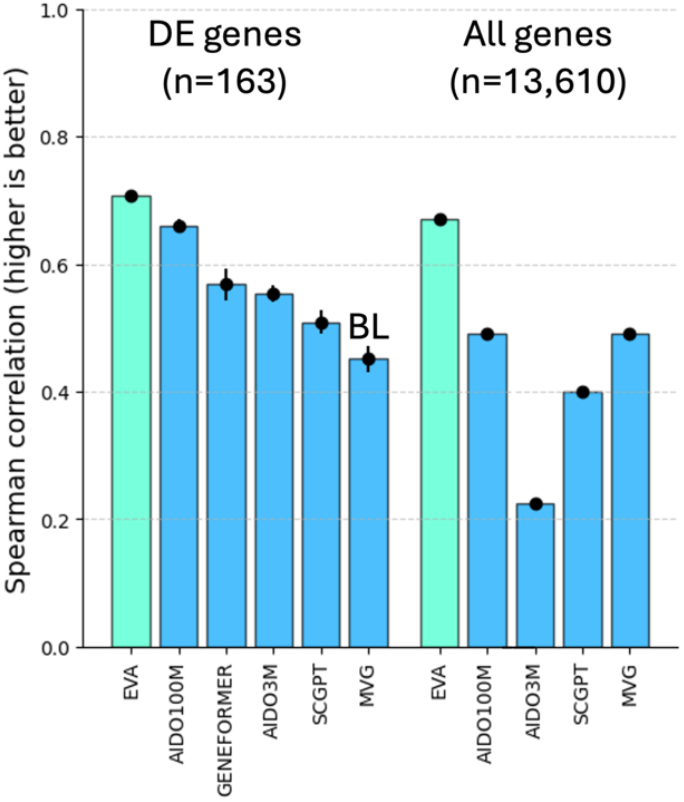
EVA learns in a few-shot setting the transcriptomic effect of TNFR2 depletion in the colon of IBD mouse models, outclassing other generalist RNA-seq foundation models. MVG is a multivariate Gaussian baseline fitted to the training data. *DE stands for significantly differentially expressed genes between perturbed and control. Results averaged over 10 random initializations; error bars mark standard deviations. (Due to its limited context size, Geneformer is only benchmarked on DE genes*.)

When focusing on the 163 differentially expressed genes between control and perturbed conditions, all foundation models outperform the Gaussian baseline model, and EVA leads the benchmark with a significant 0.70 Spearman correlation. When measuring instead the performance using all 13,610 available genes, EVA is the only robust model that continues to significantly outperform the Gaussian baseline with an almost unchanged Spearman correlation (0.68). In contrast, other foundation models fail to maintain performance, suggesting difficulty in capturing more subtle gene expression changes despite the importance of this information to characterize a coherent biological state. This comparison underscores the significance of our pre-training process on ImmunAtlas, enabling EVA to generalize more effectively to bulk IBD data compared to other RNA-seq foundation models, particularly in a few-shot learning scenario.

### EVA replicates the efficacy outcomes of an anti-TNF trial in ulcerative colitis

To further demonstrate EVA’s real-world applicability, we applied the EVA-based TNFR2 perturbation model in a realistic use case: forecasting the outcome of an anti-TNF clinical trial in ulcerative colitis (UC). Given the absence of publicly available IBD mouse datasets involving anti-TNF surrogate treatments to our best knowledge, we relied on the genetic ablation of TNFR2 as a proxy, while acknowledging that this approach may not faithfully capture the complexity of the treatment. Using UC cohort data, we created a secondary disease model using canonical machine learning and in-house observational cohort data to map patient transcriptomic state to their disease activity scores (MAYO). We then combined both models into an end-to-end drug modelling pipeline to simulate the effect of anti-TNF treatment on UC patients. In this pipeline, the change in MAYO serves as a readout, mirroring the type of outcomes typically evaluated in ulcerative colitis clinical trials.

We used the pipeline on 29 pediatric UC patient data from the PROTECT cohort^29^ with high MAYO scores (>=7) to mimic typical RCT inclusion criteria, leveraging the stochastic nature of EVA to generate 100 possible response trajectories for each of them (Fig. 3a, 3b). For each patient, we computed the expected response rate (MAYO diminution of at least 3 points or 30%) and remission rate (MAYO score lower than 2, domains were not considered) as the fraction of trajectories falling under the corresponding threshold. Given the tolerable risk, a response threshold can then be chosen (e.g. >80%). We compared the predicted population-level outcomes with those from existing anti-TNF clinical trials in IBDs^30,31^, which typically report a response rate of ∼50%. EVA predicts a median MAYO score reduction of 2.1 points (from 9 to 6.9) and a ∼44% response rate, aligning with documented anti-TNF clinical trials in UC^30,31^. However, EVA underestimates the remission rate at 0.6% compared to the expected ∼15%, possibly due to the high baseline MAYO scores, indicating strong disease markers that persist even after EVA perturbation prediction. Refining the approach by using larger cohort training data and incorporating additional biological modalities could help improving the modelling of the disease.

**Figure 3:**
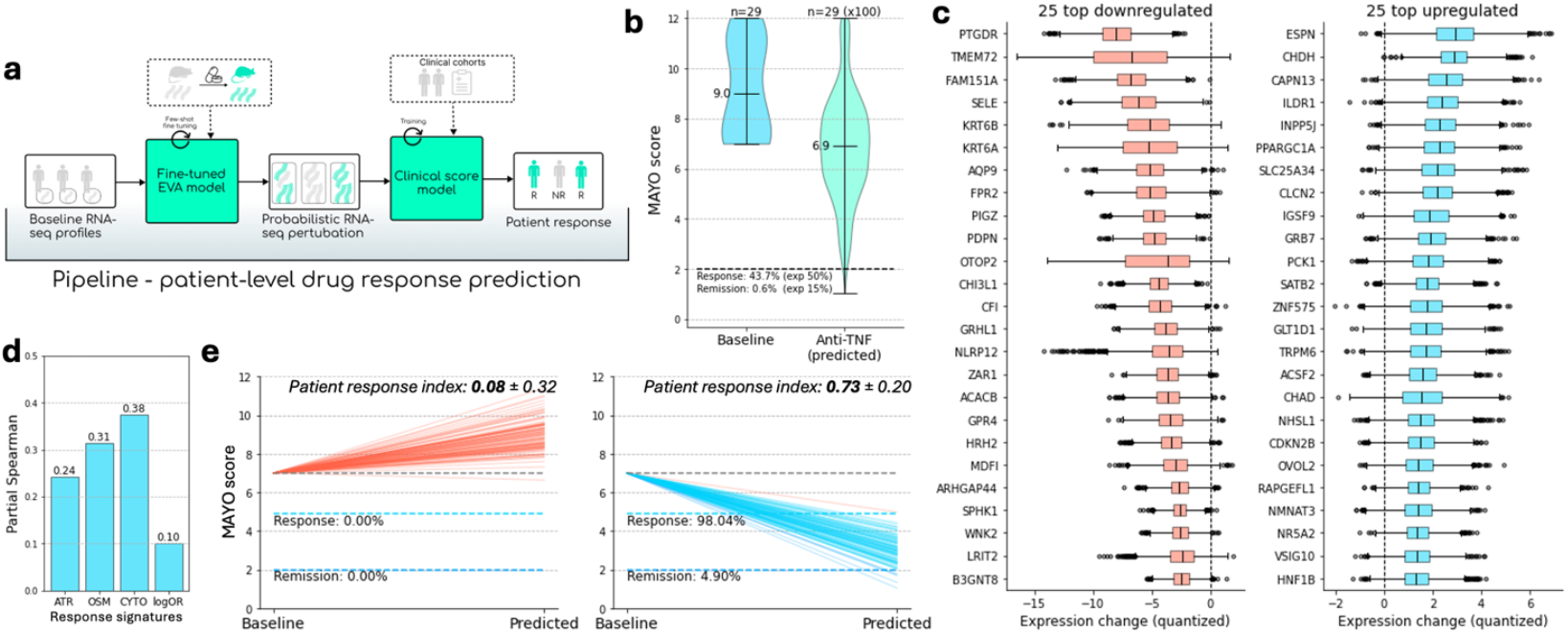
EVA replicates anti-TNF efficacy outcomes in pediatric ulcerative colitis. **(a)** Overview of the EVA pipeline. **(b)** Predicted efficacy outcomes under anti-TNF at cohort level, 100 trajectories predicted per subject, drug response measured in terms of change in MAYO score. **(c)** Top 25 downregulated and upregulated genes by EVA during the anti-TNF simulation. **(d)** Spearman correlation (corrected for baseline MAYO) between predicted response and 4 anti-TNF response signatures from the literature. **(e)** Predicted anti-TNF response for two patients with identical MAYO score at baseline. The patient response index is an aggregated z-score over the four anti-TNF signatures.

To gain insights into how EVA predicts the effects of anti-TNF treatments, we compared the expected top 25 up- and down-regulated genes with the outcomes of the inference process.

∘ Down-regulated genes include well-known inflammation-related genes (signaling: PTGDR, trafficking: SPHK1, SELE, PDPN, negative feedback: CFI, NLRP12) and others rather linked to tissue stress and renewal (KRT6A, KRT6B, OTOP2, CHI3L1, GRHL1).
∘ Conversely, up-regulated genes primarily control cell differentiation and gut homeostasis (OVOL2, SATB2, IGSF9, ESPN, ILDR1).

The model’s predictions align with expected biological responses in the inflamed gut under anti-TNF treatment, showing downregulation of both inflammation and tissue stress markers, and promotion of cell differentiation and gut homeostasis. This coherence suggests that in addition to matching the expected cohort-level response rate to anti-TNF, the model also accurately reflects the therapeutic effects of anti-TNF treatments at the molecular level in reducing inflammation and supporting tissue repair.

### EVA predicts patient-level anti-TNF response in ulcerative colitis

While forecasting realistic drug effect at the cohort level is already valuable, the ability to predict treatment response at patient level represents an even more impactful application. To evaluate patient-level predictions, we collected four transcriptomic signatures from the literature — ATR, OSM, CYTO and LogOR^32-35^— which have been associated with anti-TNF response in UC patients based on previous clinical studies. We use these transcriptomic signatures as proxies of expected patient anti-TNF response to test EVA predictions: patients with high predicted response rates should be associated with high anti-TNF response signatures. While there is still room for improvement, EVA’s patient-level predictions showed consistent correlation with all response signatures (Fig 3d), demonstrating its capability for patient-level forecasting. Despite the limited size of the inference cohorts, a finer analysis revealed cases of patients with similar baseline disease activity but radically different predicted response trajectories (two examples are shown in Fig. 3e). These differences aligned with individual signature response indices, indicating that EVA may capture biologically relevant variation not solely attributable to baseline disease severity. In this setting, EVA was able to recapitulate known patterns from a well-characterized randomized controlled trial (RCT) of anti-TNF therapy, where transcriptomic biomarkers are well established. This serves as a useful reference point to evaluate the model’s behavior. Importantly, unlike approaches that rely on predefined signatures, EVA may offer additional value in contexts such as investigational therapeutics, where such biomarkers are not yet available, by modeling patient-specific transcriptomic dependencies.

Overall, the EVA pipeline displays promising results for forecasting drug response in IMID-diseased patients. In this UC use case, its predictions align closely with the expected outcomes of clinical trials, with the additional constraints of being trained on a small dataset and focusing only a single biological modality. The consistency of the predicted response rates at the cohort level, along with the correlation between patient-level predictions and established gene expression signatures, highlights the potential of EVA for personalized medicine applications. These findings demonstrate the feasibility and scalability of leveraging large pre-trained models to simulate drug effects and predict patient responses, laying the groundwork for more targeted, effective and scalable drug development strategies in the future. Further iterations of the approach, such as the incorporation of additional biological modalities and prior knowledge, and expansion of the training dataset, will likely enhance even further the model’s accuracy and reliability.

### EVA stratifies ulcerative colitis patients and highlights potential drug response biomarkers

Beyond forecasting individual patient trajectories, the patient-level predictions generated by the EVA pipeline can be used to stratify patients based on their expected drug response, offering a powerful approach to identifying novel biomarkers. This capability is crucial for applications such as clinical trial enrichment and targeting subpopulations most likely to benefit most from a given therapy. To test this use case in our study, we applied a threshold to the predicted response rate (Fig. 4a) to categorize patients into three groups: responders (>80% response chance), non-responders (<20% response chance), and uncertain (in-between).

**Figure 4:**
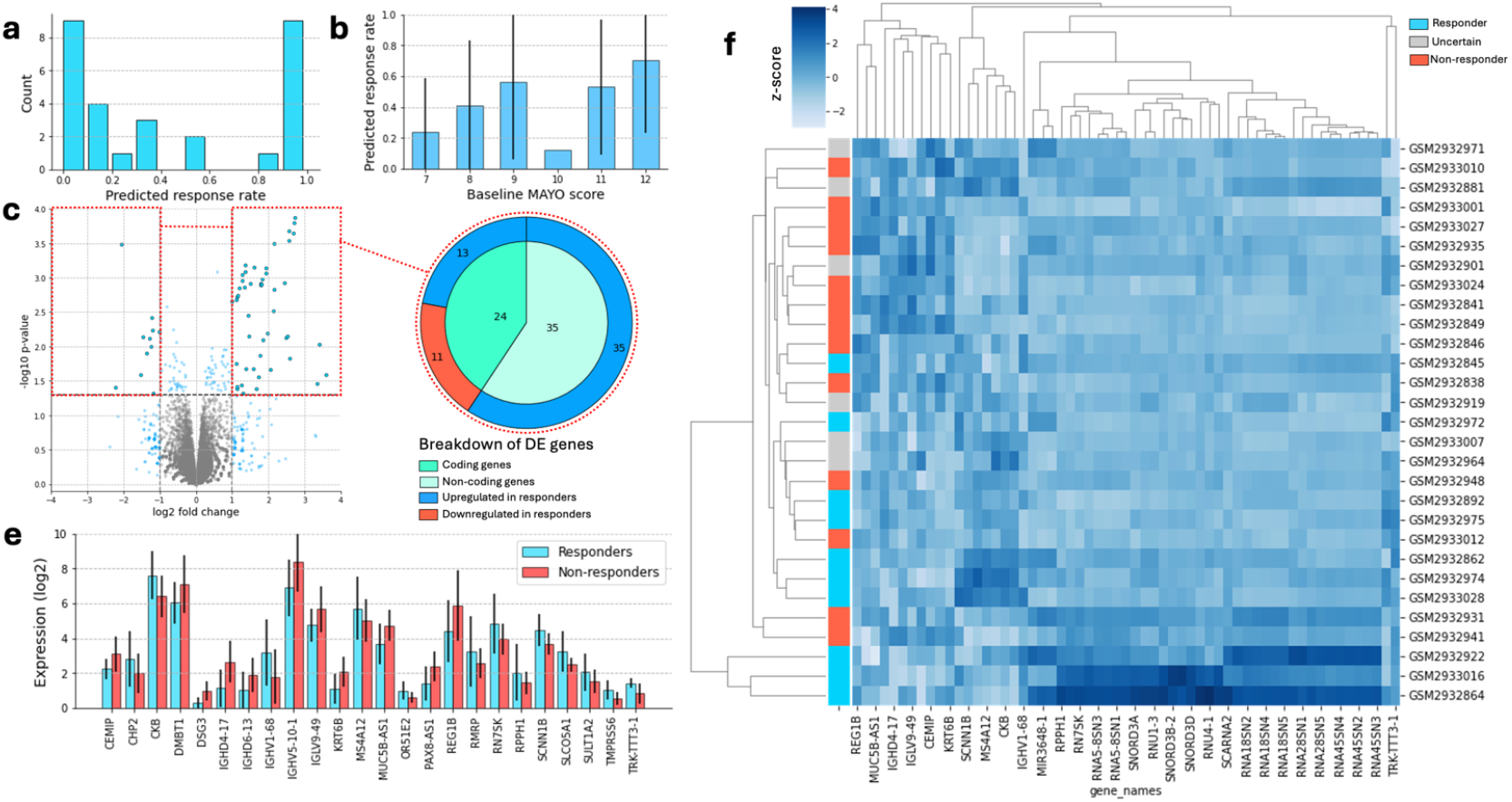
EVA enables stratification of pediatric UC patients based on predicted anti-TNF response. **(a)** Distribution of EVA-predicted anti-TNF response rates across the cohort (n=29 patients). **(b)** Average predicted response rate grouped by patients’ baseline MAYO score. Error bars represent standard deviation within each group. **(c)** Volcano plot highlighting differentially expressed (DE) genes between predicted responders and non-responders. Dashed line indicates the significance threshold. **(d)** Breakdown of significant DE genes, showing proportions of coding vs. non-coding genes and direction of regulation in predicted responders. **(e)** Comparison of expression levels (log2) for selected significant DE genes between predicted responders (green) and non-responders (red). **(f)** Heatmap displaying normalized expression (z-score) of selected DE genes (columns) across individual patients (rows).

To assess whether these predicted response groups reflect distinct biological states, we performed differential gene expression analysis between predicted high-responders and low-responders in the pediatric UC cohort. This analysis revealed numerous significantly differentially expressed genes between the two groups (Fig. 4c), encompassing both coding and non-coding genes, with varying directions of regulation (Fig. 4d). Interestingly, a significant proportion of such genes are non-coding RNAs (ncRNA), an abundant class of RNA molecules thought to play regulatory roles in molecular and cell biology^36,37^. The poor understanding of their biology usually preclude their use in knowledge-driven predictive algorithm, while foundation models like EVA could pave the way toward a better understanding of the regulatory roles exerted by ncRNA, as their expression pattern can be learned during pre-training like that of coding genes.

A heatmap of these key differentially expressed genes (Fig. 4f) reveals distinct and coherent transcriptomic signatures that characterize the predicted responder and non-responder groups. While clustering based on these selected genes is anticipated, the heatmap confirms the alignment of the stratification based on EVA’s predicted response rate with tangible differences in molecular profiles. Several groups of genes exhibit clear differential expression between the predicted groups (Fig. 4e, 4f), highlighting EVA’s understanding of specific biological pathways potentially associated with anti-TNF response variability. Remarkably, the expression of immunoglobulin segments, which could reflect tissue plasmacytosis at an advanced disease stage, but also factors indicating matrix remodeling (CEMIP, KRT6B), or stressed epithelial physiology (DSG3, MUC5B-AS1) are upregulated in non-responders. Conversely, several genes whose expression is associated with response to anti-TNF treatment are expressed in the colonic tissue under steady state. These genes are involved in varied processes of tissue homeostasis, or epithelial identity (including CHP2, CKB, SCNN1B, SLCO5A1, SULT1A2).

These findings demonstrate that EVA not only predict patient-level outcomes successfully but also uncover subgroups with distinct underlying biological states, which are relevant to treatment outcome; it further validates the pipeline’s ability to capture biologically relevant signals related to anti-TNF treatment response. It highlights EVA’s potential not only for forecasting outcomes but also for identifying molecular markers associated with treatment efficacy and stratifying patients accordingly, opening exciting opportunities for more personalized therapeutic approaches in I&I diseases.

## Discussion

In this work, we introduced EVA, a pre-trained foundation model tailored for immune-mediated inflammatory diseases (I&I diseases), to enable in-silico modelling of drug effects in a few-shot learning setting. Our study demonstrates EVA’s versatility and effectiveness in predicting drug effects, with a use case in ulcerative colitis patients using preclinical data from another species. We showcased its ability to forecast clinical trial outcomes, yielding promising results that align with expected outcomes for anti-TNF drugs, despite being trained on small datasets. This approach highlights the potential of EVA to enhance trial design and resource allocation, thereby improving the odds of success and accelerating drug development in I&I.

Beyond forecasting, EVA enables patient stratification based on predicted drug responses, offering valuable insights for personalized medicine applications. Its predictions show a strong alignment with established gene expression signatures, underscoring EVA’s potential for identifying molecular markers associated with treatment efficacy. This capability is particularly relevant for clinical trial enrichment and identifying patient subgroups most likely to benefit from targeted therapies.

This initial version of EVA comes with limitations, including its sole reliance on RNA-seq data and clinical data, which provides only partial information about the biology of I&I diseases. Future iterations will incorporate additional modalities, such as histology and proteomics data, to provide a more holistic understanding of disease mechanisms and improve predictive performance. Additionally, important challenges remain regarding the translation of findings from mouse models to human subjects and the incorporation of time and dose into the modelling process. Looking ahead, we aim to generalize the EVA framework to additional drugs and disease areas, and explore the integration of knowledge graphs, which have shown potential for significant breakthroughs when associated with foundation models, especially in a multimodal setting.

This first iteration of EVA represents a significant step forward towards a scalable, data-driven approach of drug discovery and development in I&I. We are optimistic about the future iterations and the broader implications for drug development and precision medicine.

## Acknowledgements

We’re grateful to Philippe Moingeon and Andreï Zinovyev for reviewing this work, and to all the collaborators at Scienta Lab for their contributions to this study. Special thanks to Yannis Cattan for his work on EVA pre-training, Camille Bouget for her thorough proofreading, and Jonathan Plassais for designing the graphical abstract and figures 1e and 3a. This work was granted access to the HPC resources of IDRIS under the allocation 2025-AD010316294 made by GENCI, and supported by the *France 2030* program.

## Authors contribution

AB, AF, and PM collected the data, AB and AF processed the data. AF, AB, and MC performed the software engineering and modeling. AF and MC conducted the benchmarking. PM and AF carried out the data analysis and biological interpretations. VB, JD, and PM led the study. All the authors contributed to the writing and reviewing of this manuscript and agreed to its publication.

